# High-speed volumetric fluorescein angiography (vFA) by oblique scanning laser ophthalmoscopy in mouse retina

**DOI:** 10.1101/621664

**Authors:** Weiye Song, Libo Zhou, Ji Yi

## Abstract

Oblique scanning laser ophthalmoscopy (oSLO) is a recently developed technique to provide three-dimensional volumetric fluorescence imaging in retina over a large field of view, without the need for depth sectioning. Here in the paper, we present high-speed volumetric fluorescein angiography (vFA) in mouse retina in vivo by oSLO. By simply using a low-cost industrial CMOS camera, we improved the imaging speed by ~10 times comparing to our previous results, achieving vFA at 2 volumes per second. Enabled by high-speed vFA, we visualized hemodynamics at single capillary level in 3D and provided methods to quantify capillary hematocrit, absolute capillary blood flow speed, and detection of capillary flow stagnancy and stalling. The quantitative metrics for capillary hemodynamics at 3D retinal capillary network can offer valuable insight in vision science and retinal pathologies.

## 1. Introduction

Fluorescein angiography (FA) is a major imaging method in ophthalmology, for care of retinal vascular diseases such as diabetic retinopathy (DR) and age-related macular degeneration (AMD). The fluorescein solution is administered either intravenously or orally, circulated to the retinal vasculatures, and detected upon blue light excitation.

Since the invention of FA in early 1960s (1), various retinal imaging modalities have been used to improve the resolution and image quality. Fundus photography was first used to snap-shot a two-dimensional (2D) FA from the entire field of view with a flood illumination (1). The approach is the most convenient, offering moderate resolution and image quality. Scanning laser ophthalmoscopy (SLO) applies a flying laser focus and a confocal gating to reject the diffusive signal, and significantly improves image quality (2, 3). In recent years, adaptive optics SLO (AOSLO) achieved the diffraction-limited resolution (4), and enabled FA down to individual capillaries with high-speed (5-7). However, all the above modalities primarily present FA as 2D images, either integrating signals from different retinal layers or from a particular depth section. The 2D presentation of start-of-the-art FA is a major disadvantage for a three-dimensional (3D) retinal vasculature.

To overcome the disadvantage, we recently developed a novel imaging method, named oblique scanning laser ophthalmoscopy (oSLO), and demonstrated *in vivo* volumetric FA (vFA) over 30° viewing angle with cellular resolution in 3D (8, 9). By using an off-axis illumination, an oblique light sheet can be generated by a scanning laser (8-10). At the same time, an oblique imaging system is aligned such that a camera sensor is conjugated with the oblique light sheet to capture a cross-sectional FA image (8-12). This mechanism uniquely enabled vFA by only one raster scan, without the need for depth sectioning. Our previous *in vivo* demonstration further adopted a scanning protocol for optical coherence tomography angiography (OCTA) for multimodal imaging(8). Herein in this paper, for the first time to our knowledge, we demonstrated *in vivo* vFA imaging on mouse retina by oSLO with significantly increased the imaging speed, achieving vFA at 2 volumes per second (vps). The improved high-speed oSLO allows us to capture and quantify hemodynamics at individual capillary level in 3D.

## 2. OSLO image notation

The unique feature of oSLO is to capture cross sectional fluorescent images in retina, analogues to the perspective in OCT. We therefore follow the similar image notation. The 3D vFA image is in *x, y, z* axis, where *x* and *y* is the fast and slow scanning directions in the transverse/lateral plane, respectively; and *z* is the depth dimension. One cross sectional frame in *x-z* by oSLO is a B-scan, consisting of numeral A-lines.

## 3. System and Imaging Methods

### 3.1 System setup

The system setup is modified from our previous publication [8]. The visible light (λ<650nm) was filtered out by a dichroic mirror (DM1) from a super-continuum laser source (Superk EXTEME, NKT Photonics). The beam then was polarized, and dispersed by a pair of identical prisms (P1, P2). A beam block passed the blue light from ~420nm to 490nm. The beam was reflected and picked by a D-shaped mirror, then collimated to a single mode fiber. The fiber delivers the blue light to an *f* = 6 mm collimator (L1), relayed by two sets of telescope systems (L2:L3, L4:L5) to eye pupil, and steered by two galvanometer scanning mirrors (GM1, GM2). The second 3:1 telescope system (L4:L5) is mounted on a customized dove tail slider, which offsets the optical axis to create the oblique scanning illumination [8]. The shift of the dove tail slider is ~3mm. The power of blue excitation on the pupil is 30 μW. The fluorescence emission from the eye was reflected to the detection optical path by a dichroic mirror (DM3) placed in the middle of the 3:1 telescope. Another telescope (L5:L6) relays the emission light from the pupil plane to the de-scanned galvanometer mirror (GM3). Then the beam is relayed to the objective lens (OL2, Olympus UplanFL N 20 ×/0.5) by another 1:1 telescope (L7:L8). The final imaging system (OL3, Olympus UplanFL N 10 ×/0.3, and L9, Navitar F2.8/50mm) is mounted on a 3 axis stage (X, Y, and angle) to capture images by a high-speed CMOS camera (BFS-U3-51S5M-C, PointGray, Canada). All the fluorescence detection optics system is mounted on a 2 axis translational stage and the angel between final imaging system and optical axis is ~30 degrees.

**FIGURE 1.**
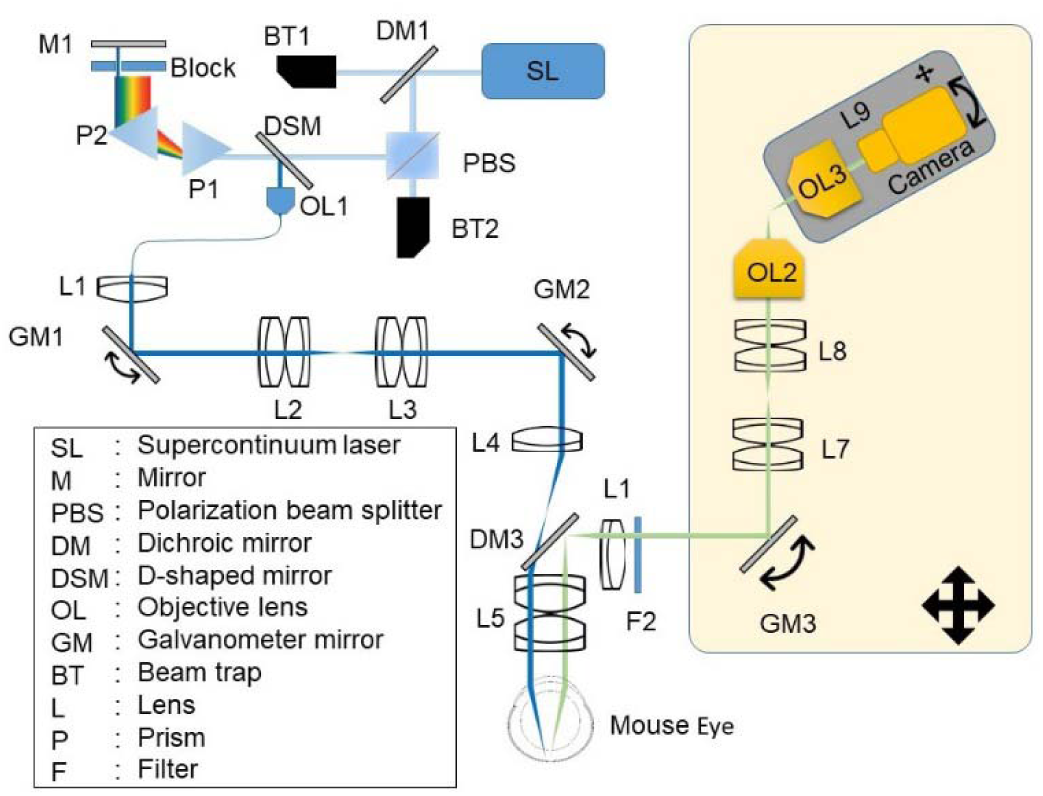
Schematic of the oblique scanning laser ophthalmoscopy (oSLO). The component within the shaded area are mounted on a 2-axis translational stage.

### 3.2 Image acquisition protocol

The scanning protocol is modified from an OCTA scanning protocol. Briefly, an 80% duty cycle saw tooth pattern was sent by an analog output board (NI, PCIe-6731) to control the fast *x* scanning galvanometer mirror (GM1). A ramping voltage was sent for GM2 for slow *y* scanning. The period of the saw tooth is 2.5 ms. The saw tooth pattern was repeated twice for each *y* scanning step for one B-scan. The exposure time of the camera is 5 ms. To maximize the speed, we used 2×2 binning and output 1224×300 pixels for B-scan in *x,* and *z*. The frame rate is limited to 200 Hz due to the overhead in readout from the camera. The scanning density and field-of-view (FOV) is controlled by a Labview software for different experimental protocols. For high-density and high-speed vFA, 512 and 100 B-scans were acquired along *y* direction, respectively. To continuously monitor the hemodynamics at individual capillary segments, we repeatedly performed B-scan at one *y* location over time. To calculate the velocity of capillary blood flow, a high-density vFA scan was first performed to map vessels in 3D, and immediately followed by alternating B-scans at two closely spaced *y* locations (alternating B-scans) for 300 times at a 200Hz frame rate.

**Table 1.**
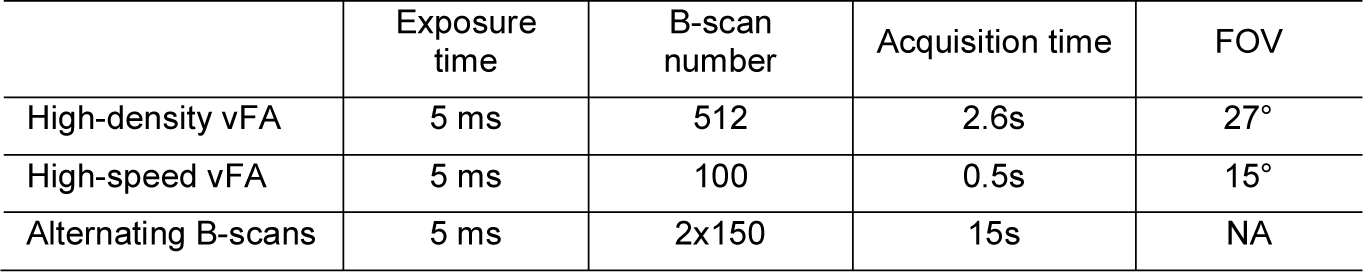
Summary of different vFA imaging protocols

### 3.3 Image processing

In order to analyze vFA images at different vascular layers, we referenced the depth to outer plexiform layer (OPL) where the deep capillary plexus is located. We manually pinpointed ~10 capillaries at OPL within the cross sectional vFA images every 10 B-scans. Within those selected frames, a 2^nd^ polynomial fitting was applied to the coordinates (*x, z*) of the annotated capillaries to identify OPL. Then another 2^nd^ polynomial fitting was applied on the coordinates (*y, z*) of OPL along slow scanning direction for the entire volumetric dataset. By referencing to OPL, vFA signal at different retinal layers can be obtained by segmenting the signal from a desired depth range. While the eye movement under anesthesia do not significantly impact high-speed imaging, the repeated vFA dataset can share the same OPL reference. Otherwise, the image processing would be required for each dataset.

### 3.4 Animal preparation

All procedures were approved by the Institutional Animal Care and Use Committee of Boston University Medical Center. Wild-type C57BL/6J mice (male at 8 weeks, Jackson Laboratory) were used in this study. Anesthetized mice introduced by intraperitoneal injection of Ketamine (80 mg/kg) and Xylazine (10 mg/kg) was placed on a custom-made 5-axis (x, y, z translations, yaw and pitch) holder to allow adjustments of eye position and angle. 0.1 ml 10% FITC-dextran (Sigma-Aldrich, mol wt 40,000) was injected intravenously. 1% Tropicamide ophthalmic solution was applied to the mice’s eye to dilate the pupil for 2 minutes first, then commercial artificial tears was applied to the mice’s eye to keep it moist during the experiment. A thermal lamp was used to provide heating during the experiment. After imaging, the animal was released and placed in a recovery box. The animal was closely monitored until it has regained sufficient consciousness to maintain sternal recumbency.

## 4. Results

### 4.1 Volumetric fluorescein angiography (vFA) in mouse retina in vivo

Figure 2a exemplifies a high-density vFA from a mouse retina *in vivo*. By using a CMOS camera, we improved the imaging speed by ~10 times from our previous publication (8, 9), and one high-density vFA can be acquired in ~2.5 seconds over a 27° angle of view. Two cross sectional vFA images clearly show the distribution of microvasculature through the retina to choroid. We averaged vFA signal within a region of interest (ROI), and plotted it against the depth axis (Fig 2b). There appears four major peaks, indicating retinal nerve fiber layer (rNFL), inner plexiform layer (INL), outer plexiform layer (ONL), and choroid. The distance from rNFL to OPL is roughly equal to the one from OPL to choroid, consistent with the mouse retina anatomy that the inner and outer retina has roughly the same thickness.

The 3D capability of vFA allows us to segment and display individual microvascular layer in rNFL, IPL, OPL and choroid, as shown in Fig. 2c-2f. The vascular branching can be observed from the major arterioles in rNFL, to intermedia plexus in OPL, and finally the capillary plexus in OPL. One interesting feature is that some capillaries appear merging to venules in OPL, and directly back to major vein (Fig. 2e), consistent to the retinal vascular structures observed in other reports in mice (13, 14).

**FIGURE 2.**
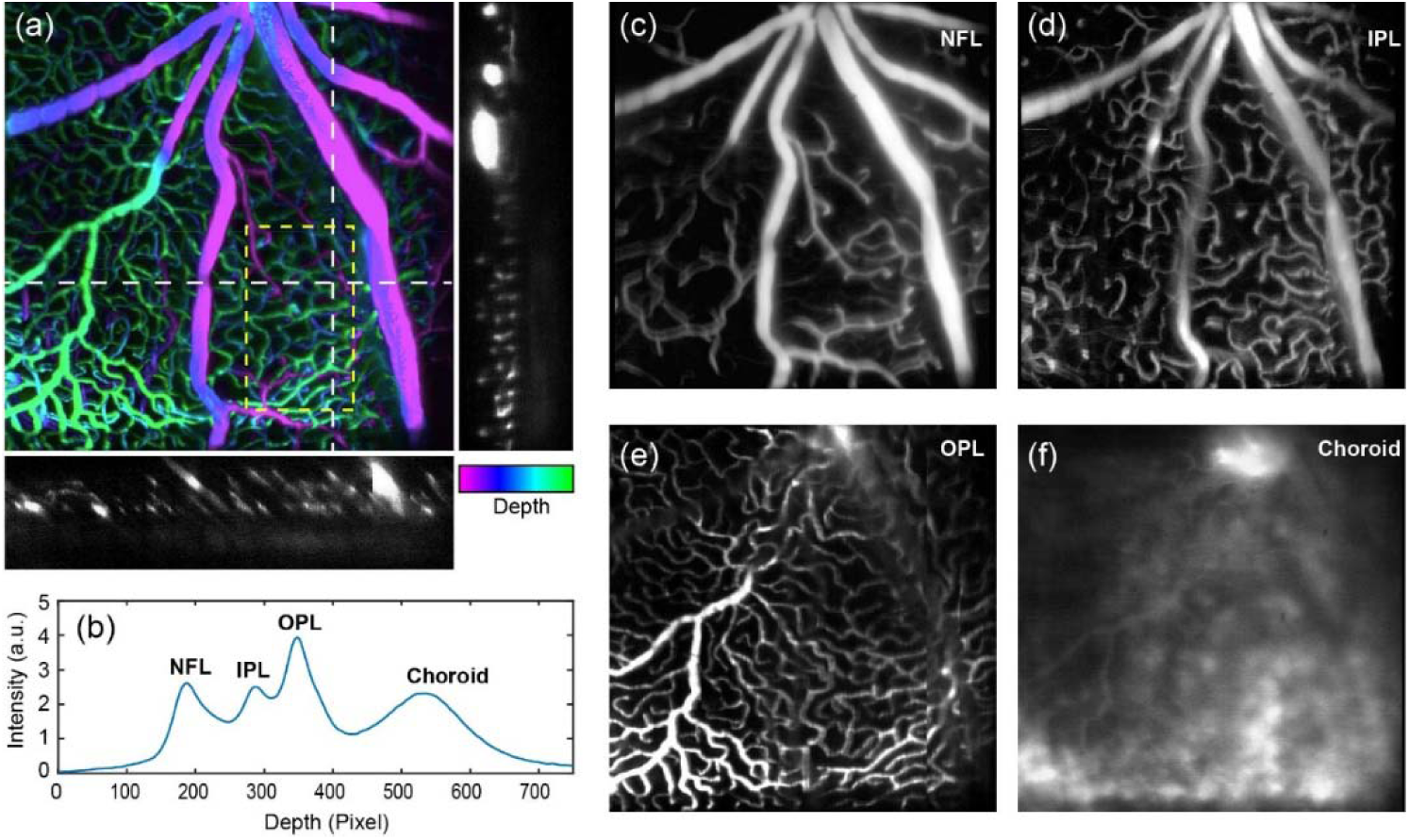
Volumetric fluorescein angiography (vFA) in mouse retina *in vivo*. (a) A depth-encoded vFA image from a mouse retina over a 27° field-of-view. The pseudo-color is in HSV space. The depth coordinate of the maximum intensity denotes the hue, and the image intensity denotes the saturation and value. Two cross-sectional images are exemplified along the white dash lines. (b) The depth distribution of the vFA signal within the yellow region of interest in panel (a). NFL: nerve fiber layer; IPL: inner plexiform layer; OPL: outer plexiform layer. (c-f) The averaged vFA from NFL, IPL, OPL, and choroid, respectively. The contrast was adjusted separately for each layer.

### 4.2 Capillary hematocrit via temporal averaging

The high-molecular weight FITC-dextran is confined within an intact retinal microvasculature and does not diffuse intracellularly. When we repeatedly acquire B-scans at one *y* location, the temporal FA signal would exhibit an intermittent pattern at a single capillary when blood cells pass in a single file, where bright and dark signal denote plasma and blood cells respectively (Fig. 3a) (7). Figure 3b demonstrated one experimental results. The 3D resolving capability of vFA allows high confidence of locating capillaries in OPL, and the intermittent fluorescence signals can be visualized at 200Hz B-scan frame rate.

**FIGURE 3.**
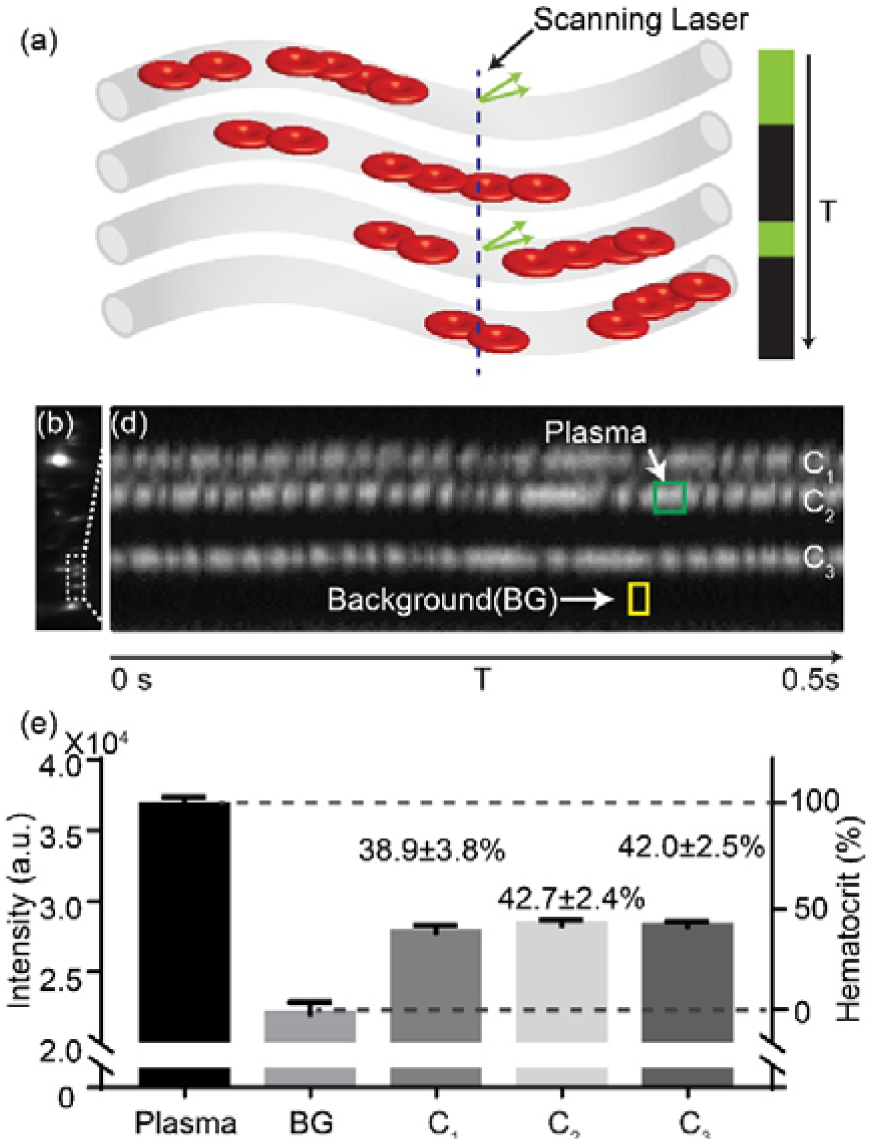
Capillary hematocrit calculation via temporal averaging. (a) Illustration of the fluorescence signal generation with repeated B-scans at a single capillary. T is short for time. (b) An example of a cross-sectional vFA image from a mouse retina *in vivo.* (c) The temporal vFA signal from three capillaries (C1, C2, and C3) in OPL from panel (b). The green and yellow regions exemplified the region-of-interests (ROIs) for calculating vFA at 0% hematocrit in plasma, and virtually 100% hematocrit in a non-vascular area. (d) The quantitative hematocrit calculation based on the vFA signal intensity with temporal averaging every 0.5s over a total period of 2.5s. For plasma and BG, five ROIs were averaged (n=5). For C1 to C3, five measurements were averaged over a total 2.5s data (n=5). Bar = SEM.

We first calculated hematocrit (Hct, %), defined by volume percentage of blood cells in whole blood. It was recognized that bright blocks in Fig. 3b were from plasma with 0% hematocrit, and the non-vascular area (BG) could equivalently and virtually represent 100% hematocrit when no FITC-dextran was present. We then randomly selected five large bright blocks, and averaged the image intensity of center regions (Fig. 3c) to quantify the vFA signal at 0% Hct, as *I*_0%Hct_. Similarly, the mean image intensity of five non-vascular areas quantifies the vFA signal at 100% Hct, as *I*_100% Hct._ After temporally averaging the vFA signals *I*, Hct at any capillaries can be quantified by

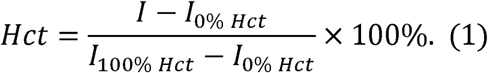

Figure 3d shows the results of Hct calculation from the three capillaries shown in Fig. 3c. The mean Hct values were around 40%, close to the value under the normal physiological conditions. Hct from C1 to C3 were calculated every 0.5 second, over a total period of 2.5s. The error bar represents the temporal Hct variation.

### 4.3 Absolute capillary blood flow speed

The cluster and spacing between blood cells at individual capillaries give rise to the intermittent temporal pattern in vFA signals, as demonstrated in Fig. 3. When sampled at two closely spaced points from a capillary segment, two vFA temporal patterns are likely similar without branching or merging. The absolute capillary blood flow speed can be deterministically calculated, given the time delay between two temporal patterns and the distance between the two sampling points.

To implement the concept, we modified our scanning protocol from Fig. 3, such that a high-density vFA is first taken, immediately followed by alternative B-scans at two closely-spaced *y* locations for 3 seconds. The high-density vFA was to locate the capillary segments in 3D. Since each B-scan took 5ms, the sampling rate for the two alternating B-scans was 100 Hz. Figure 4a shows the overview of the deep capillary plexus at OPL segmented from the high-density vFA, proceeding to the alternating B-scans. The intersected capillaries can then be easily identified. The temporal patterns at two locations of the same capillary (C_1_ and C_2_) are plotted in Fig. 4b, with similar patterns and a visible time delay. We calculated the temporal cross-correlation for the time delay of *ΔT* [s] (Fig. 4c), and the blood speed *v* [m/s] was calculated by

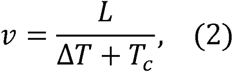

where *Tc* [s] is the time for each B-scan, 5 ms in our case, and *L* is the Euclidean distance between C_1_ and C_2_ that was calculated from the 3D high-density vFA. Note that we used OPL as the depth reference, and the dimension herein is calculated in 2D. Importantly, the sign of *ΔT* can be either positive or negative, indicating the flow direction.

**FIGURE 4.**
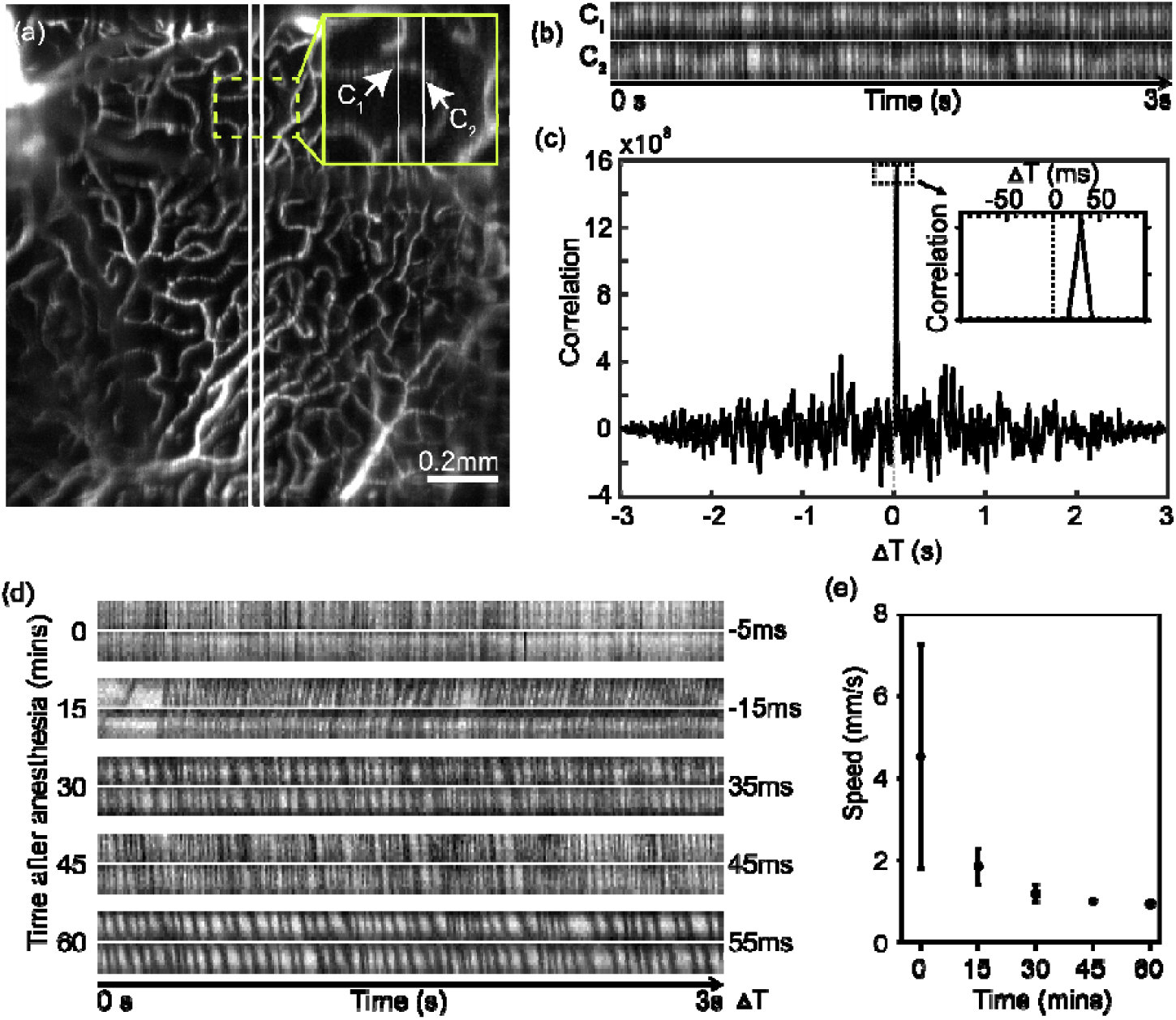
Measurements of absolute blood speed at individual capillary level. (a) vFA image at deep capillary plexus at outer plexiform layer (OPL). Two alternating B-scan locations are labeled with two white lines. Bar = 0.2 mm. (b) The temporal vFA signal at two capillaries, C_1_ and C_2_, as pointed out in panel (a). (c) The temporal correlation of two temporal vFA signals between C1 and C2. The peak of the correlation measures the time delay. (d) Examples of vFA temporal signal from five capillaries, with various blood velocity. (e) The capillary flow speed measured longitudinally from the same retina after anesthesia (n=5). Bar = SEM.

To demonstrate the capability of longitudinally measuring capillary blood speed, we imaged a normal C57BL/6 mouse retina every 15 minutes after the Ketamine/Xylazine anesthesia over a course of 60 minutes. The capillary in OPL was separated to measure the blood velocity, and the temporal signal pair from capillaries at each time point are exemplified in Fig. 4d. While the pattern may be inconsistent sporadically, the overall correlation and time delay were rather clear. The frequency of the vFA signal intermittence decreases and the time delay increases, both indicating a slower blood speed at capillary level after prolonged anesthesia. The statistical result was shown in Fig. 4e, the mean velocity of the capillary in OPL was ~4.5 mm/s immediately after anesthesia and dropped to ~0.9 mm/s 60 minutes later. This reduction may be due to the body temperature drop and therefore reduced blood pressure over the course of the experiment.

### 4.4 Quantitative metrics for 3D retinal capillary hemodynamics

At a 200 Hz frame rate, we achieved 2 vps vFA with 100 B-scans per volume. Figure 5 shows the capillary plexuses located in IPL and OPL at different time stamps separately, during a course of 15s acquisition. The intermittent vFA signals within capillaries are also visible due to the passing blood cells (See Visualization 1).

**FIGURE 5.**
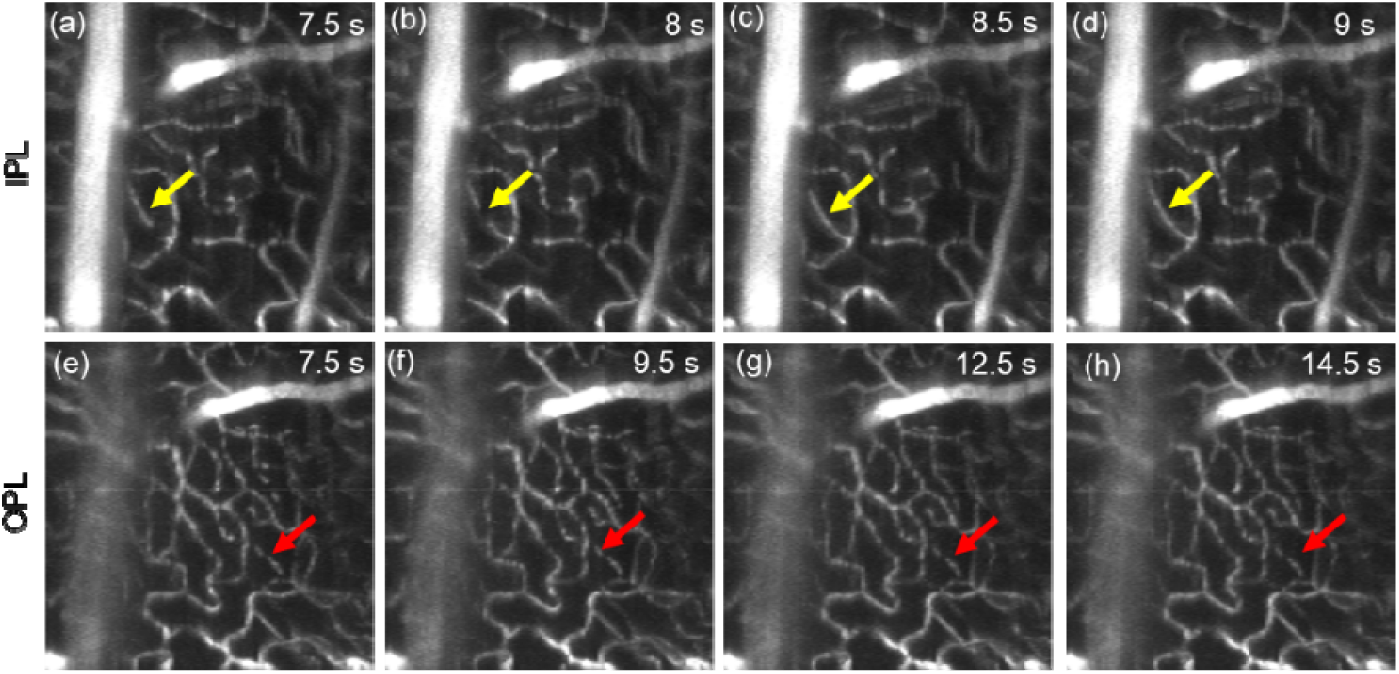
Capillary flow dynamics in high-speed vFA at 2 volume per second segmented at the inner plexiform layer, IPL (a-d) and outer plexiform layer, OPL (e-h). Yellow arrows point to one capillary segment showing a transient ischemia, and the red arrows point to a capillary having stalled blood flow.

We used two metrics: hematocrit, and frame-to-frame image cross-correlation, to quantitatively characterize the capillary hemodynamics in 3D vFA. For hematocrit, the similar approach was taken as described in Fig. 3, such that we took vFA intensity from a bright segment of capillary and the non-vascular background to represent 0% and 100% Hct, respectively. We then manually segmented each capillary, and the mean value of the vFA intensity with the segment quantified Hct by Equ. 1. Fifteen capillaries from IPL and OPL were segmented (Fig. 6a, 6b), and a heat map of hematocrit with summarized the selected segments over time. While the mean hematocrit is ~40% (Fig. 6d) over 15s, the temporal variation for each capillary, as well as the spatial variation among different capillaries are apparent from the heat map. For segment #3, there appears a transient ischemia for 1s, followed by a quick recovery, which is easily appreciated in Visualization 1.

**FIGURE 6.**
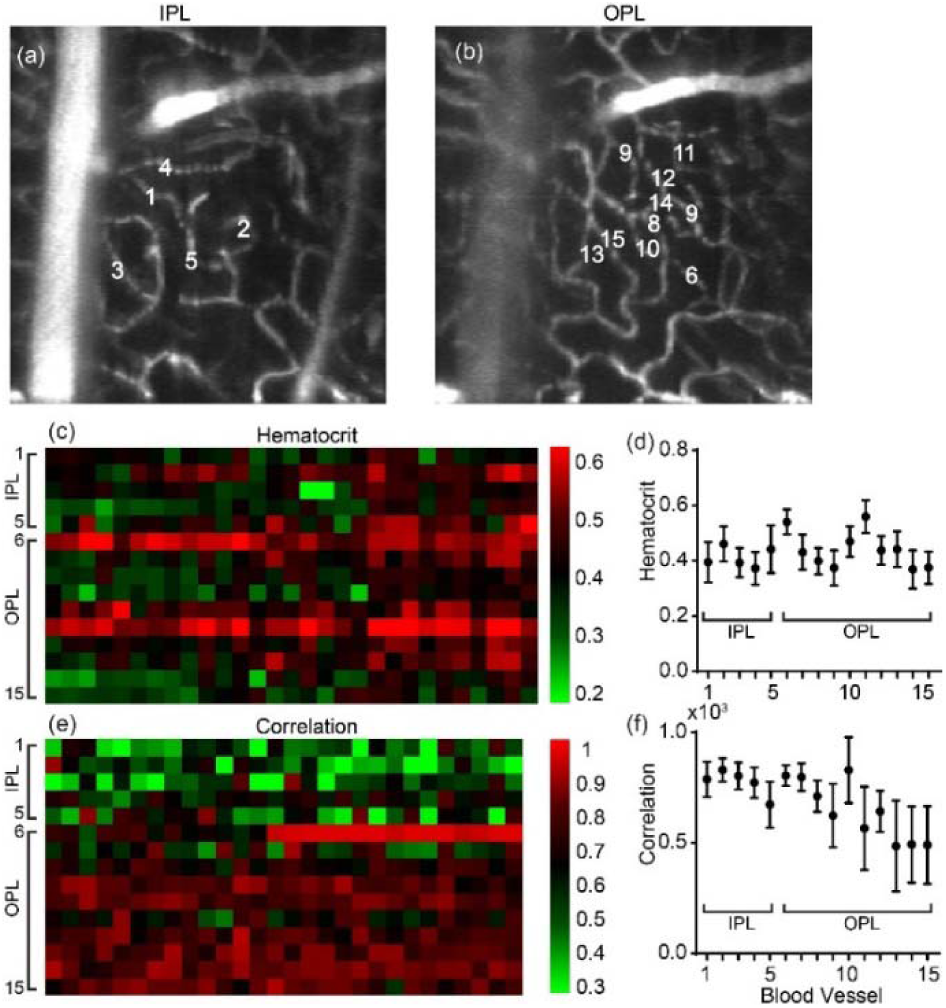
(a-b) vFA segmented at inner plexiform (IPL) and outer plexiform layer (OPL) with denoted capillary segments. (c)The heat map showing longitudinal hematocrit change at each capillary segment over a total period of 15s with 0.5s interval. (d) The averaged hematocrit over 15s for each capillary segment labeled in panel (a-b) (n = 15). (e) The heat map showing the temporal image correlation between two vFA collected 0.5s apart within each capillary segment. (f) The averaged image correlation averaged over a total period of 15s (n = 14). Bar = SEM.

Within each segment, we then calculated the image cross-correlation between adjacent frames to detect the capillary blood flow stagnancy and stalling as the stagnancy would result in high image cross-correlation. A heat map was also generated to present the temporal and spatial variation (Fig. 6e). Capillary stalling can be observed in segment #6 with absence of blood flow for several seconds (See Visualization 1). We can also observe overall higher cross-correlation within OPL than IPL, suggesting slower blood flow in the deep capillary plexus presumably to allow better oxygen perfusion (Fig. 6e-6f).

## 5. Discussion

Herein we presented a high-speed volumetric fluorescein angiography (vFA) by oblique scanning laser ophthalmoscopy in mouse retina *in vivo*. For the first time to our knowledge, 3D fluorescein distribution within mouse retinal microvasculature at capillary level *in vivo* can be rapidly imaged at 2 vps. The imaging speed is significantly improved over our previous results in rat retina (8, 9), and the high speed volumetric imaging allows measurement and quantification of capillary hemodynamics. We further provided quantitative metrics to measure capillary hematocrit and absolute capillary blood flow speed.

Using intermittent fluorescein signal for capillary hemodynamics has been previously reported (7). The advantage of using oSLO is that we now can discern capillary FA signals from different depths at a cross-sectional view, without the need for any depth sectioning. The 3D capability allows segmentation of specific retinal layers excluding the choroidal background, and the confounding signals from other inner retinal layers. This leads to high confidence of locating individual capillary segment from a relatively large field of view, as demonstrated in Fig. 4. Without the capability of depth segmentation, the temporal correlation methods at two adjacent B-scan location will be confounded by signals from overlapping capillaries in different depth.

The imaging speed for vFA ultimately defines the highest capillary blood flow speed measurable in Fig. 4. In Equ. 2, the least increment of the temporal correlation *ΔT* is 2×5=10ms without signal interpolation, at the speed of 200 Hz frame rate. The estimation of *L* is ~36 μm in our alternating B-scan protocol, and thus the maximum projective speed *v*_p_ is ~7.2mm/s. This detection range appears to suffice at the normal physiological conditions for mice that the average velocity in Fig. 4e was 4.5 mm/s immediately after anesthesia. The detection range can also be expanded by increasing the distance between the two alternating B-scans. The same consideration is applied in longitudinal vFA imaging in Fig. 5. At this 2 vps, cautions need to be applied to interpret the results, since vFA image temporal correlation in Fig. 6e-6f does not directly measures blood flow speed particularly when the flow is fast. However, when the capillary flow is significantly stalled or slowed down, the temporal correlation should be correlated to the blood speed, as can be seen for capillary segment #10 in Fig. 6e.

The limitation of the current oSLO setup is the reduced image quality at the peripheral image at ~30° viewing angle. This is largely inherent with the aberration in mouse eye, which may be effectively reduced using contact lens (15). The imaging speed can also be improved by using more sensitive detector, and optimization of the imaging system.

High-speed vFA provides unique opportunities to characterize the hemodynamics at the capillary level over a 3D volume of view. Hematocrit, blood flow speed, and capillary stalling provides a comprehensive view of the perfusion function within the microvascular system, which is both fundamentally and pathologically important in broad range of diseases, such as cancers, neurodegeneration, diabetic retinopathy, and macular degeneration. The quantitative metrics can be valuable approach for the purpose of diagnosis and phenotyping within a living biological system.

## Funding

National Institute of Health (NIH) (R01NS108464, R21EY029412, R01CA224911, R01CA232015); Bright Focus Foundation (G2017077, M2018132).

## Disclosures

Ji Yi held patent for oblique scanning laser ophthalmoscopy, and may have financial interest in the commercialization of oSLO. Other authors declare no conflicts of interest related to this article.

## Notes

https://www.youtube.com/watch?v=Jye_wshQrms

